# Gasdermin D is activated but does not drive neurodegeneration in SOD1^G93A^ model of ALS: Implications for targeting pyroptosis

**DOI:** 10.1101/2024.10.10.617609

**Authors:** Georgia Gunner, Himanish Basu, Yueting Lu, Matthew Bergstresser, Dylan Neel, So Yoen Choi, Isaac M. Chiu

**Affiliations:** Harvard Medical School, Department of Immunology, Blavatnik Institute Boston, MA 02115

## Abstract

Amyotrophic Lateral Sclerosis (ALS) is a fatal neurodegenerative disease characterized by progressive motor neuron loss, microgliosis, and neuroinflammation. While pyroptosis, an inflammatory form of programmed cell death, has been implicated in ALS, the specific role of Gasdermin D (GSDMD) – the primary executioner of pyroptosis – remains unexplored. In this study, we examined the function of GSDMD in the well-established SOD1^G93A^ mouse model of ALS. Our results showed robust GSDMD activation in the spinal cords of SOD1^G93A^ animals across two strain backgrounds, with elevated expression in Iba1+ microglia. To explore its role in disease progression, we bred B6.SOD1^G93A^ mice onto a GSDMD*-*deficient background. In comparing SOD1^G93A^; *Gsdmd*+/+ and SOD1^G93A^; *Gsdmd*−/− mice, we found that *Gsdmd* loss did not affect disease onset, weight loss, or grip strength decline in either male or female animals. Notably, GSDMD deficiency resulted in a modest but statistically significant increase in mortality in SOD1^G93A^ mice. Moreover, GSDMD absence had minimal impact on astrogliosis, microgliosis and motor neuron loss. These findings show that while GSDMD is activated in the ALS mouse model, its loss does not mitigate key ALS behavioral phenotypes, gliosis or motor neuron loss. This study provides insights into the potential therapeutic relevance of targeting pyroptosis and inflammatory pathways in ALS.

## 1. Introduction

Amyotrophic lateral sclerosis (ALS) is a fatal neurodegenerative disease characterized by progressive loss of motor neurons in the spinal cord, accompanied by both central and peripheral neuroinflammation^1,2^. Mutations in the superoxide dismutase (SOD1) gene are a leading cause of familial ALS^3^, inducing toxic misfolding and aggregation of SOD1 protein, which disrupts cellular homeostasis in neurons and glial cells^4^. The SOD1^G93A^ mouse model faithfully recapitulates many facets of ALS in human patients, including progressive motor neuron loss, microglia activation, astrogliosis, and paralysis, with mortality around six months of age^5,6^. It is the most commonly used preclinical model of motor neuron disease.

Microglia, the resident myeloid cells of the CNS, are essential for synapse pruning and maintenance of neuronal health. However, their aberrant activation can drive neurodegeneration. Innate immune responses in microglia and other non-neuronal cells have been shown to play important roles in regulating motor neuron health in ALS and disease progression^7^. Pro-inflammatory cytokine production and microglial expansion are prominent features of ALS, both in mouse models and postmortem human tissues^8^.

Emerging evidence implicates the pyroptosis pathway in ALS and other neurodegenerative diseases including Alzheimer’s disease, Parkinson’s disease, multiple sclerosis, and traumatic brain injury^9^. Pyroptosis is an inflammatory cell death pathway activated by the inflammasome complex and caspase-1/caspase-11 cleavage^10,11^. Gasdermin-D (GSDMD), the primary executioner of pyroptosis, is expressed in macrophages and microglia in an autoinhibited form. Upon inflammasome activation, caspase-1 and/or caspase-11 cleaves GSDMD, releasing its N-terminal domain, which subsequently targets the plasma membrane to oligomerize and form large pores^12–14^. These GSDMD pores facilitate the release of pro-inflammatory cytokines, such as IL-1 and IL-18, and eventually lead to cell membrane lysis via NINJ1^15,16^. In some cases, GSDMD pores can form on viable cells, such as bone marrow-derived macrophages (BMDMs), resulting in a hyperactivated state without cell death^17^.

In ALS patients, spinal cord and motor cortex tissues display increased expression of NLRP3, GSDMD, and IL-18, and components of the NLRP3 inflammasome as well as caspase-1 activation are evident in resident glia of the CNS^18^. The SOD1^G93A^ mouse model similarly shows elevated expression of GSDMD, caspase-1, NLRP3, and Il1B in the lumbar spinal cord^19^. Cultured primary microglia from SOD1^G93A^ mice show increased expression of components of the inflammasome including NLRP3, IPAF, AIM2, and ASC^20^. Addition of exogenous mutant SOD1 to mouse microglia also drives IL-1B cytokine release and caspase 1 activation^21–23^. Moreover, caspase-1 and IL-1beta deficiency has been shown to increase lifespan, reduce gliosis, and improve motor neuron survival in SOD1^G93A^ mice^21^. These data indicate a potentially major role for pyroptosis activation and downstream IL-1 release in driving ALS disease progression.

Currently, the activation of the inflammasome is a clinical target in neurodegeneration. Specifically, NLRP3 antagonists are being pursued to treat neurodegeneration in Alzheimer’s disease, Parkinson’s Disease, ALS, and other disease conditions^24,25^. Contrary to the effect of blocking caspase-1 or IL-1B, studies using the NLRP3 inhibitor MCC950 showed that chemical inflammasome inhibition does not extend survival or reduce cell death in the SOD1 G93A mouse model or in iPSC-derived motor neurons^26,27^. Thus, further study is warranted to elucidate the role of pyroptosis in preclinical models of ALS.

We hypothesized that GSDMD, as the key executioner of pyroptosis that allows for IL-1 release, may mediate neuroinflammation and pathology in ALS. To test this hypothesis, we used the SOD1^G93A^ transgenic mouse model of ALS, which overexpresses mutant human Cu/Zn superoxide dismutase^28,29^. Microglia were identified as the main GSDMD-expressing cell type during disease progression. We observed that GSDMD activation and cleavage increased with disease onset, correlating with caspase-1 activation in the spinal cord. To assess the functional significance of GSDMD, we generated SOD1 G93A mice lacking GSDMD (SOD1^G93A^; *Gsdmd−/−*). While GSDMD deletion did not significantly impact disease onset, grip strength, or motor neuron loss, it modestly but significantly accelerated mortality. These findings suggest that although GSDMD is activated in SOD1G93A spinal cords, its loss does not ameliorate disease pathology. Thus, targeting GSDMD should be approached with caution as therapeutic strategy for ALS.

## 2. Materials and Methods

### 2.1 Animals

All mouse experimental procedures were performed in compliance with the Harvard Medical School and The Jackson Laboratory Institutional Animal Care and Use Committees. C57BL/6NJ (JAX # 005304), C57BL/6N-Gsdmd^em4Fcw^/J (JAX # 032410), and SOD1^G93A^ mice (JAX # 002299) mice were obtained from Jackson labs and bred at Harvard Medical School. Animal experiments were fully approved by the Harvard Medical School Institutional Animal Care and Use Committee (IACUC). Animals were housed in temperature (22 ± 2 °C) and humidity (55 ± 5%) controlled care facilities at Harvard Medical School on a 12 h light:dark cycle and provided with freely available food and water.

The SOD1^G93A^; *Gsdmd −/−* mice were obtained by crossing male mice carrying the SOD1^G93A^ transgene with female *Gsdmd −/−* (JAX # 032410) mice for 3 generations. The control SOD1^G93A^; *Gsdmd +/+* mice were obtained by crossing SOD1^G93A^ mice with wild type female mice having a B6NJ background (JAX #005304), also for 3 generations. This breeding strategy controlled for any strain background related phenotypic differences. All offspring were tested for equivalent SOD1 gene copy numbers by qPCR (forward primer: CAGTAACTGAGAGTTTACCCTTTGGT; and reverse primer: CACACTAATGCTCTGGGAAGAAAGA). Mice that had SOD1 signal drop-off of more than 30% from control SOD1G93A mice were considered as low copy number animals and were excluded from the study. Both male and female mice were used for all experiments. Animals were provided with food and water *ad libitum*.

### 2.2 Western Blotting

Tissue lysis: For analysis of mouse spinal cords, animals were anesthetized with Avertin solution (500 mg kg−1, MilliporeSigma) and perfused with 20 ml of cold PBS before harvest. Spinal cords were dissected in dish of cold PBS and stored at −80 C prior to tissue homogenization/lysis. Lysis buffer was prepared on ice as follows: 10 ml T-per buffer (Thermo #78510), 1 tablet protease inhibitor (Sigma #11836153001/Roche), 100ul HALT protease inhibitor (Thermo #87786), 100ul 0.5M EDTA (Thermo #87786), 1 tablet Phosstop™-phosphatase inhibitor tablets (Sigma #04906845001/ Roche). For tissue lysis, frozen spinal cords were resuspended in cold lysis buffer in a 2mL Eppendorf tube. One autoclaved metal bead (BB) was placed in each tube containing sample and lysed in a bead beater (Qiagen TissueLyser II) at a setting of 25 Hz for 5 minutes. Homogenized tissue samples were then incubated on a rotating tube rack at 4C for 30 min. Following incubation samples were clarified by spinning at 16000 × g for 15mins at 4C. Supernatants (lysate) were then taken and stored at −80 C prior to analysis. For immunoblot processing, sample lysates were diluted in Bolt 4X loading buffer (Thermo #B0007) containing 1X Bolt Sample Reducing Agent (Thermo #B0009) and beta-mercaptoethanol. Samples were then boiled for 10 min at 90 C, spun down and stored at −20 C.

Immunoblotting: Samples were incubated on 90C heat block for 10 minutes and run on Bolt 4–12% Bis-Tris-Plus gels (Thermo Scientific cat# NW04125BOX). Gels were transferred to nitrocellulose iBlot 2 membranes (Fisher Scientific cat# IB23001), blocked with 5% Pierce Clear Milk Blocking Buffer (Thermo Scientific cat# 37587) for 30 minutes, washed 3x with TBST (TBS, 0.05% Tween-20), and incubated overnight in blocking buffer containing primary antibody at 4C. Primary antibodies included rabbit anti-GSDMD (Abcam cat# ab219800, 1:800), rabbit anti-Caspase 1 (AdipoGen cat# AG-20B-0042-C100, 1:500), rabbit anti-Iba1 (Cell Signaling Technology cat# 17198) and mouse anti-GAPDH (EMD Millipore, cat# CB1001, 1:1000). Following primary antibody incubation, blots were washed 3x with TBST for 10 min, incubated with secondary antibody for 1h at RT, followed by three additional 10 min washes with TBST. GSDMD and Iba1 immunoblots (secondary antibody rabbit anti-HRP Cell Signaling Technology cat # 7074S, 1:1000) were developed with Supersignal West Pico Chemiluminescent Substrate (Thermo Scientific cat# 34080) on a Azure 300 chemiluminescent imaging system. GAPDH and Caspase-1 immunoblots were incubated with IR-fluorophore conjugated secondary antibodies (LI-COR Biosciences cat# 926–32213, cat #926-32210, 1:10,000) and developed and imaged on a Li-COR imaging system.

### 2.3 Immunohistochemistry

Adult mice were perfused with 20mL of cold PBS and 20mL of cold 4% PFA. Spinal cords were dissected, fixed overnight in 4% PFA at 4C, paraffin embedded and processed for IHC. Paraffin embedded sections were dehydrated in successive washes with xylene and ethanol. Sections were then washed in water and boiled in 1X Citrate Buffer unmasking solution (Cell Signaling cat# 14746S). Tissue sections were then incubated in a 3% H_2_0_2_ solution for 10 min, washed in TBST-Tween20 and blocked for 1h at room temperature with TBST/5% Normal Goat Serum (Cell Signaling Cat# 5425). Following blocking, tissue sections were incubated in diluted primary antibody solutions (see below) overnight at 4C. The next day, sections were washed with 1X TBST and incubated for 30min at room temperature in SignalStain® Boost IHC detection Reagent (HRP rabbit, #8114 or HRP mouse, #8125) specific to the species of the primary antibody. Slides were washed with TBST prior to Tyramide Signal Amplification (TSA). Fluorescein conjugated TSA reagent (Akoya Biosciences, NEL745001KT) or Cy3 reagent (Akoya Biosciences, NEL744001KT) were diluted as per manufacturer’s instructions. Slides were incubated with TSA reagent for 10min at room temperature (protected from light). Slides were then washed three times with 1X TBST, counterstained with DAPI and mounted in prolong gold antifade mounting medium (Cell Signaling cat# 8961S) and imaged using a Leica Thunder microscope with Andor Zyla sCMOS camera via a 20× Plan Apo objective. For serial staining of sections (dual IHC) a stripping step was performed by boiling slides for 10 min in a 10mM sodium citrate buffer (Cell Signaling cat# 14746). Following stripping, slides were the incubated in primary antibody solution, washed, incubated with SignalStain Boost IHC secondary detection reagent and subject to TSA amplification as described above. Primary antibodies used: rabbit anti-GSDMD (Abcam, cat# ab209845, 1:800), rabbit anti-Iba1 (Cell Signaling Technology cat# 17198), rabbit anti-GFAP (Sigma Aldrich, cat # G3893-100UL, 1:1000), Green Fluorescent Nissl Stain (Invitrogen, cat# N21480, 1:1000).

### 2.4 Mining of published RNAseq databases

GSDMD expression was performed in RStudio from the single cell RNAseq of healthy mouse spinal cords published by Blum et al 2021 (PMID: 33589834).

GSDMD expression in microglia and sciatic nerve macrophages of the spinal cords of SOD1G93A mice was performed in RStudio from the published database published by Chiot et al 2020 (PMID 33077946).

### 2.5 SOD1^G93A^ behavioral analysis

Weight Measurement: Weights of all mice carrying the SOD1^G93A^ transgene were measured biweekly from week 7 to week 21. Disease onset was defined as the day of peak weight while disease progression was defined as the number of days between disease onset to euthanasia.

Grip strength: Grip strengths of the SOD1^G93A^ transgenic mice were measured weekly, starting at week 7 and ending at week 21. A grip strength machine (BIOSEB# bio-GS3) with a mesh grid attachment was used to obtain combined grip strengths of all four limbs. For each measurement, the mouse was held at the base of the tail and placed on the mesh grid. A total of 5 pulls were performed per mouse with approximately 5–10 second breaks between each pull. The top three pulls per time point were averaged and used for data analysis. 12 or more mice per genotype, per sex, were used for weights and grip strength analysis.

Survival: Each mouse was monitored daily after symptom onset. Mice that were unable to rear were given hydrogels and wetted food pellets at the bottom of the cage. Euthanasia time points for each mouse were determined by the inability to successfully right itself when flipped on either of its sides within 30s.

### 2.6 Quantification and Statistical Analysis

All statistical analysis was performed in GraphPad Prism (version 10.0.1, GraphPad Software, USA). Data are reported and plotted a Mean +/− Standard Error of the Mean (SEM). Statistical significance was determined using a two-tailed unpaired Student’s t-test for normally distributed data. One-way or two-way ANOVAs were used for comparisons involving more than two groups. For multiple comparisons, a post hoc analysis using Tukey’s or Šídák’s test was performed to determine differences between groups. For SOD1 survival and disease onset data, Kaplan-Meier curves were constructed, and survival analysis was performed. Groups were compared for statistical differences using the log-rank test (Mantel-Cox). Results were considered statistically significant where P < 0.05.

## 3. Results

### 3.1 GSDMD is activated in SOD1^G93A^ spinal cords with disease progression

The SOD1^G93A^ transgenic mouse is a well-established model that recapitulates the progressive paralysis and mortality seen in human ALS patients^28,29^. Disease progression in this model varies depending on the genetic background, with the mixed B6/SJL SOD1^G93A^ transgenic mice showing a faster progression compared to B6 congenic SOD1^G93A^ mice^30^.

We analyzed GSDMD protein levels over time in spinal cord homogenates from SOD1^G93A^ or non-transgenic (nTg) control mice on the two different genetic background strains. At baseline, GSDMD exists in its full-length form (53 kDa), and upon activation and cleavage by caspase-1, the N-terminal fragment (30 kDa) is generated. By western blot analysis, SOD1^G93A^ mouse on the C57Bl/6NJ (B6) background showed significantly increased levels of the N-terminal, activated GSDMD form compared to control non-trangenic (nTg) controls at both day 140 (mid-stage) and day 162 (end-stage) of disease (genotype, F_(1, 7)_ = 6.236, P=0.0412) (Figure 1 A, C). Full-length GSDMD also increased, indicating that expression of GSDMD itself may also increase over time in addition to its activation (Figure 1 B). GSDMD cleavage correlated with increased expression of both full length caspase-1 (age, F_(1, 7)_ = 8.578, p =0.0221; genotype F_(1, 7)_ = 25.61, p = 0.0015) and activated caspase-1 (20kDa) (genotype, F_(1, 7)_ = 102.8, p <0.0001) Figure 1 A, D-E). We also analyzed mice harboring the SOD1^G93A^ mutation on the mixed B6/SJL background (B6.SJL). In this strain, onset of disease is significantly earlier than SOD1^G93A^ transgenic mice on a pure C57Bl/6J background^30^. At presymptomatic disease age of d60, little cleaved N-terminal GSDMD is evident in the whole spinal cord protein lysate in both nTg and SOD1^G93A^ mice (Figure 1 F). However, at day 120 (end-stage) of disease, an increase in both full-length and cleaved N-terminal GSDMD is evident in B6SL.SOD1^G93A^ mice (Figure 1 G-H). Original, uncropped blots are shown in Supplemental Figure 1.

**Figure 1:**
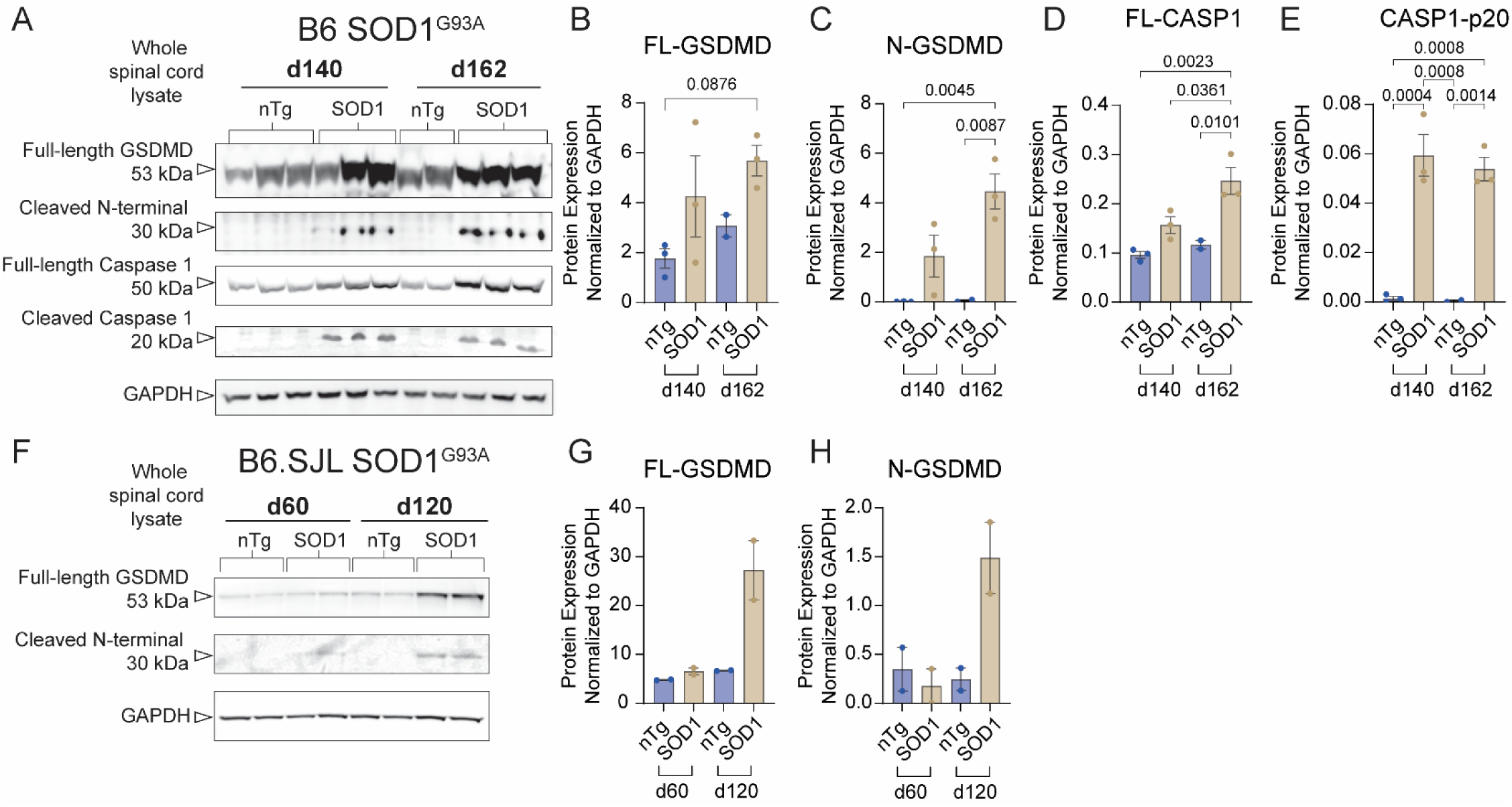
GSDMD is activated in SOD1-G93A mouse spinal cords on both B6 and B6.SJL backgrounds. (A) Western blot of whole spinal cord protein lysate of C57Bl/6NJ SOD1^G93A^ animals at 140 days (mid-stage disease) and 162 days (end stage). N = 3 nTg d140 animals, n = 3 SOD1 d140 animals, n = 2 nTg d162 animals, n = 3 SOD1 d162 animals. (B-C) Quantification for full length GSDMD (FL-GSDMD) (B) and cleaved N-terminal GSDMD (N-GSDMD) (C) bands normalized to loading control GAPDH. (D-E) Quantification of full-length caspase-1 (FL-CASP1) (D) and cleaved caspase-1 (cCASP1) species p20 (E) normalized to GAPDH. (F) Western blot of whole spinal cord protein lysate from C57Bl/6-SJL SOD1^G93A^ animals at 60 days (presymptomatic) and d120 (end-stage). N = 2 nTg d60 animals, n = 2 SOD1 d60 animals, n = 2 nTg d120 animals, n = 2 d120 SOD1 animals. (G-H) Quantification of full length GSDMD (FL-GSDMD) (G) and cleaved N-GSDMD (H) normalized to GADPH. Statistics for (B-E) was performed using a Two-Way ANOVA with Tukey’s multiple comparisons test.

### 3.2 GSDMD is primarily expressed by microglia and increases in expression in symptomatic SOD1^G93A^ animals

We next asked where GSDMD is expressed in the spinal cords of wild-type non-transgenic (nTg) and SOD1^G93A^ mice at non-symptomatic (d100), early (d120) and mid-stage disease (d140). We found via immunohistochemistry (IHC) that gasdermin-D (GSDMD) expression is minimal but highly localized to microglia labeled with Iba1 in nTg control mice at day 140 of age (Figure 2A). To confirm this expression pattern of GSDMD in the spinal cord, we mined the published single cell RNAseq database of resident spinal cord cells in healthy, adult mice from Blum *et al* 2021^31^ (Supplemental Figure 2 A). Analysis of this dataset found that GSDMD is expressed in two cell types in the spinal cord: microglia and endothelial cells. IHC performed in the spinal cords of SOD1^G93A^ mice at mid-stage disease (140 days), revealed GSDMD expression is highly increased but still localized to Iba1+ cells (Figure 2 B). Overall quantification of GSDMD immunostaining showed that total GSDMD positive area in the lumbar region of the spinal cord increased with disease progression from d100, d120 to d140 (F_(2, 15)_ = 22.01, p<0.001) (Figure 2 C).

**Figure 2.**
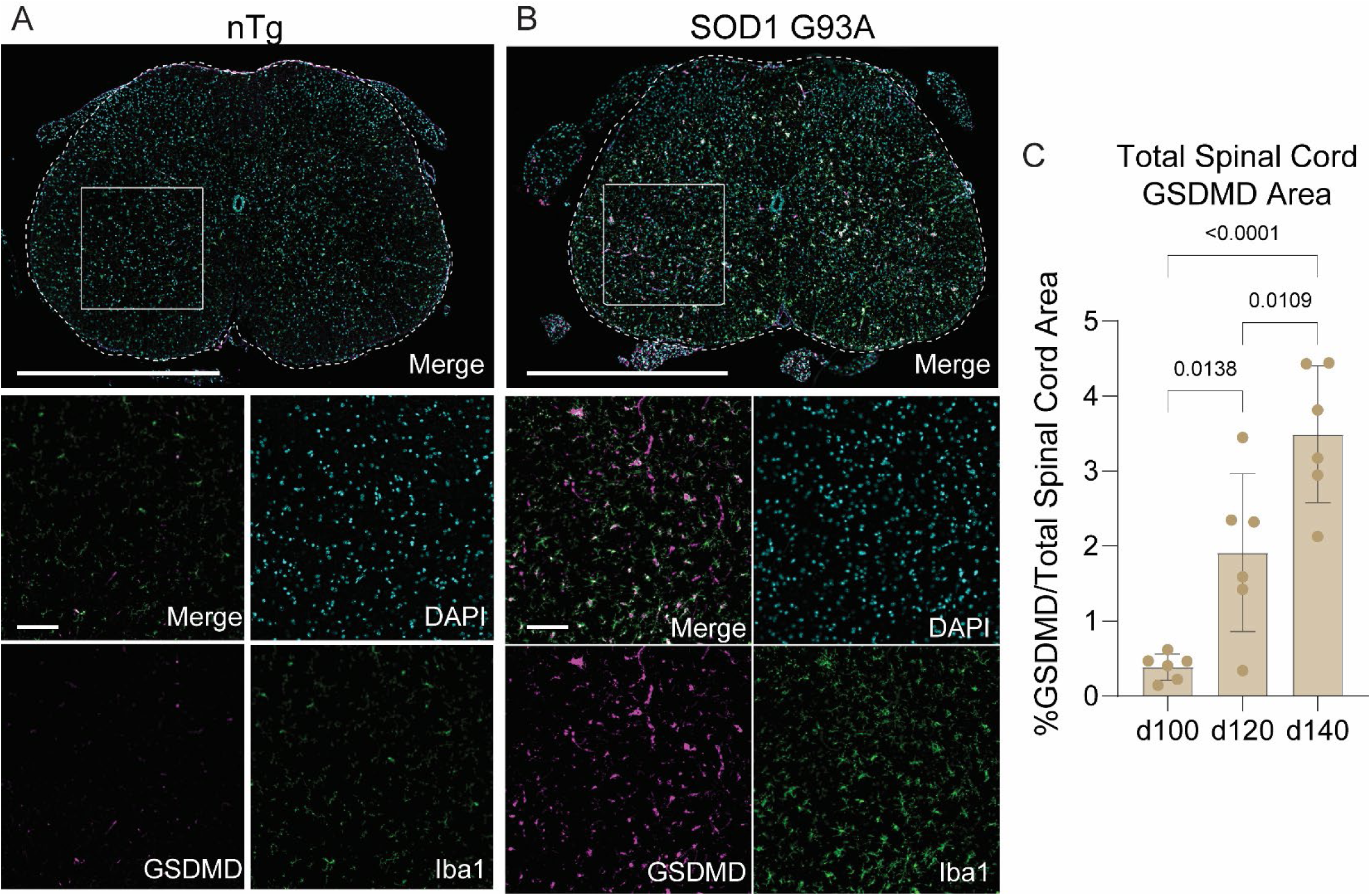
GSDMD immunostaining increases in ventral horns of SOD1^G93A^ mouse spinal cords. (A) Representative image of lumbar spinal cord of nontransgenic (nTg) C57Bl/6NJ mouse at 140 days stained with microglia marker Iba1 (green), GSDMD (magenta) and nuclei (DAPI, cyan). Scale bar = 1000 um. Inset images scale bar = 100 um. (B) Representative image of lumbar spinal cord of C57Bl/6NJ SOD1G93A mouse at 140 days stained with markers listed in (A). Scale bar = 1000 um. Inset images scale bar = 100 um. (C) Quantification of total GSDMD area over entire spinal cord area over time. N = 3 spinal cord sections averaged per N = 6 SOD1G93A animals. Statistics for C performed using One-Way ANOVA with Tukey’s multiple comparisons test.

We next analyzed the degree of GSDMD found in Iba1+ microglia and whether it changed in intensity or localization over the course of disease progression by performing immunohistochemistry (Figure 3 A-C). Using Iba1 to label microglia in the spinal cord, the percent of GSDMD co-localized to Iba1+ cells significantly increased as animals age (F_(2, 15)_ = 21.50, p<0.001) (Figure 3 D). The fluorescence intensity of GSDMD signal internalized within microglia also significantly increased with aging (F_(2, 15)_ = 15.71, p<0.0002) (Figure 3 E). While we see GSDMD is highly co-localized to resident microglia, we know from scRNAseq databases of the spinal cord that endothelial cells also express *Gsdmd* (Supplemental Figure 2). In fact, if we quantify the total level of GSDMD signal that is colocalized to Iba1+ microglia at each age, we see that by d140 there is a significant decrease in the overall GSDMD signal that is co-localized to microglia (F_(2, 14)_ = 4.064), p = 0.0406) (Figure 3 F). This could be attributable to an upregulation in GSDMD expression in endothelial cells, but it could also be possible that other resident CNS cells, such as astrocytes, could also increase GSDMD expression in the context of mutant SOD1-driven cell dysfunction.

**Figure 3.**
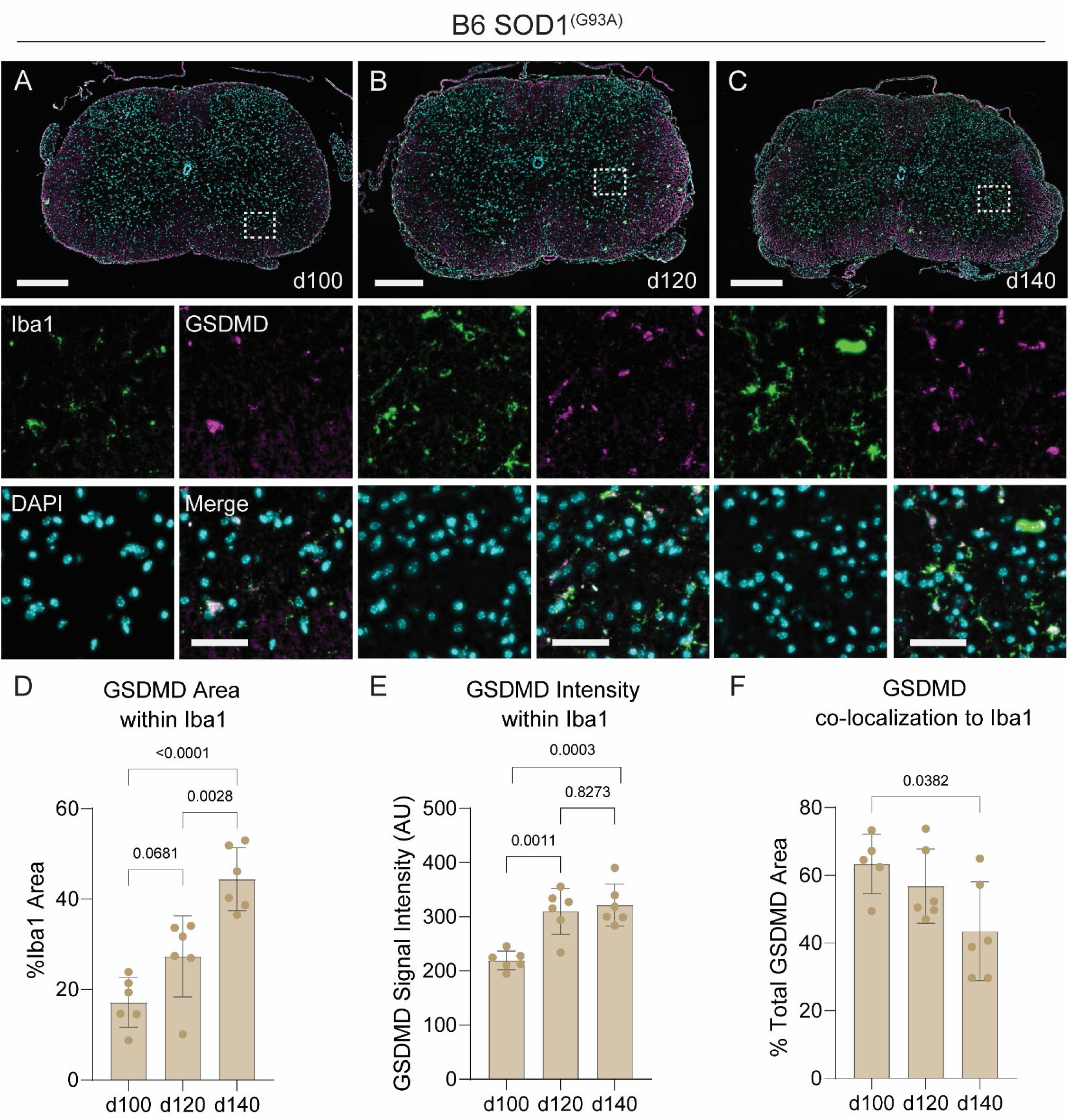
GSDMD is upregulated in Iba1+ microglia in SOD1^G93A^ spinal cord over time. Representative images of B6 SOD1^G93A^ lumbar spinal cords used for quantification at pre-symptomatic (100 day) (A), early disease (120 day) (B) and mid-stage (140 day) disease (C). Sections stained with microglia marker Iba1 (green), GSDMD (magenta) and nuclei marker DAPI (blue). Scale bar = 250um. Insets focus on ventral horn region. Scale bar = 50 um. (D) Quantification of total GSDMD area co-localized (within) to Iba1+ cells over time. (E) Quantification of GSDMD signal intensity within microglia over time. (F) Quantification of the percent of total GSDMD area co-localized to Iba1 over time. (D-F) N = 3 spinal cord sections averaged per N = 6 SOD1^G93A^ animals. Statistics for performed using a One-Way ANOVA with Tukey’s multiple comparisons test.

To determine whether GSDMD expression changes at the transcriptional level over time in myeloid cells in SOD1^G93A^ mice, we mined published sequencing databases of SOD1^G93A^ sciatic nerve macrophages and spinal cord microglia^32^ (Supplemental Figure 2 B). From this database, we found SOD1^G93A^ microglia upregulate *Gsdmd* expression at the onset of disease with higher expression as animals age (cell type F_(1, 29)_ = 14.32, p = 0.0007). Peripheral sciatic nerve macrophages also highly expressed *Gsdmd* in SOD1^G93A^ mice, though levels in these cells did not change greatly with time (Supplemental Figure 2 B). The increase in GSDMD at both the protein level via IHC and evidence for transcriptional upregulation from published RNAseq data led to our hypothesis that microglial GSDMD expression may be critical in driving disease severity in the SOD1^G93A^ model.

### 3.3 SOD1^G93A^ transgene expression in WT and *Gsdmd* deficient background

To generate a genetic model of null *Gsdmd* expression on the SOD1^G93A^ transgenic mouse background, we set up a cross of *Gsdmd* −/− animals to the SOD1 G93A (B6) model. As the *Gsdmd* −/− line is on a C57Bl/6NJ background, we crossed both wild-type C57Bl/6NJ and *Gsdmd* −/− to the SOD1 G93A B6 line for 3 generations. This backcross led us to generate SOD1^G93A^; *Gsdmd−/−* transgenic mice with SOD1^G93A^; *Gsdmd+/+* controls with similar background expression of the C57Bl/6NJ inbred strain. We first determined that our cohort of animals for characterization of the disease model harbored a similar degree of SOD1-G93A transgene overexpression (Figure 4 A), as it has been well established that copy number of the mutant SOD1 transgene significantly changes the rate and severity of paralysis onset^33^ (sex, F_(1, 70)_ = 0.4639, p = 0.4981; genotype, F_(1, 70)_ = 0.04716, p = 0.8287; interaction, F_(1, 70)_ = 2.969, p = 0.0893). To validate complete knockout of *Gsdmd* expression, we also performed immunohistochemistry for GSDMD in the spinal cord of both SOD1^G93A^; *Gsdmd−/−* and SOD1^G93A^; *Gsdmd+/+* mice (Figure 5 A, B). SOD1^G93A^; *Gsdmd−/−* display no expression for GSDMD, which we quantified focusing on the ventral horn (t=4.499, df=7, p = 0.0028) (Figure 5 B). In addition, we isolated CD11b+ cells from the spinal cord of SOD1^G93A^; *Gsdmd−/−* and SOD1^G93A^; *Gsdmd+/+* mice at d140 (mid-stage disease) and performed a Western blot for GSDMD protein to additionally confirm loss of GSDMD expression (Figure 5 C). Original, uncropped blots are shown in Supplemental Figure 3.

**Figure 4.**
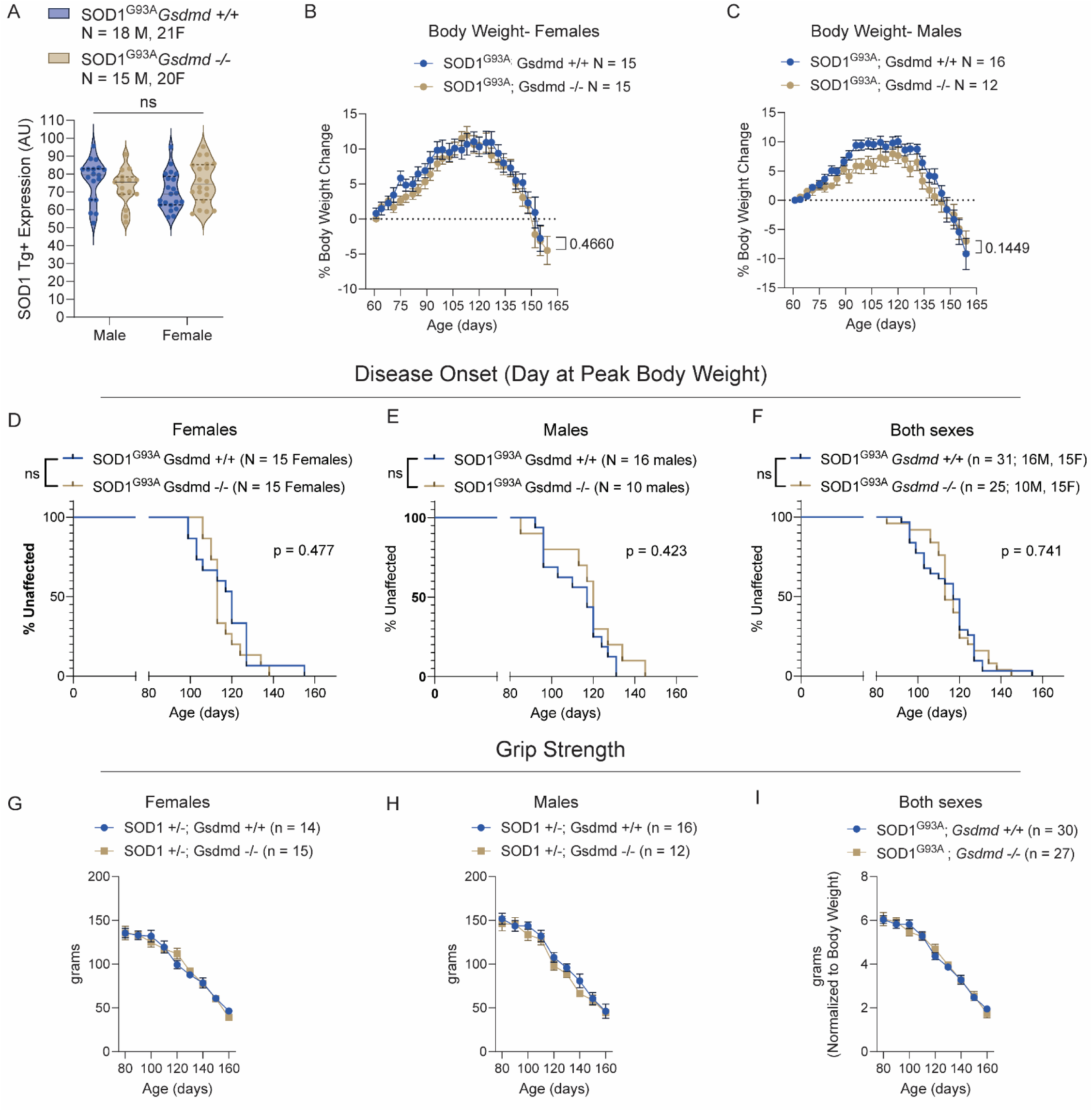
SOD1^G93A^; *Gsdmd* −/− animals display no difference in loss of body weight, disease onset, or grip strength compared to SOD1^G93A^; *Gsdmd* +/+ animals. (A) Quantification of SOD1 transgene overexpression in the cohort of male and female SOD1^G93A^; *Gsdmd*+/+ and SOD1^G93A^; *Gsdmd* −/− animals used for phenotyping and survival. Statistics performed using a Two-Way ANOVA with Tukey’s multiple comparisons test. (B-C) Quantification for the percent body weight change from baseline at day 60 for female (B) and male (C) SOD1^G93A^; *Gsdmd*+/+ and SOD1^G93A^; *Gsdmd* −/− animals. Statistics performed using a Two-Way ANOVA with Šídák’s multiple comparisons test. (D-F) Quantification for disease onset, defined as the age at which animals reach peak body weight, for female (D), male (E) and combined sexes (F) for SOD1^G93A^; *Gsdmd*+/+ and SOD1^G93A^; *Gsdmd* −/− animals. Statistics performed using a Log-rank (Mantel-Cox) test. (G-I) Grip strength measured weekly for female (G), male (H) and combined sexes (I) for SOD1^G93A^; *Gsdmd*+/+ and SOD1^G93A^; *Gsdmd* −/− animals. Statistics performed using a Two-Way ANOVA with Šídák’s multiple comparisons test.

**Figure 5.**
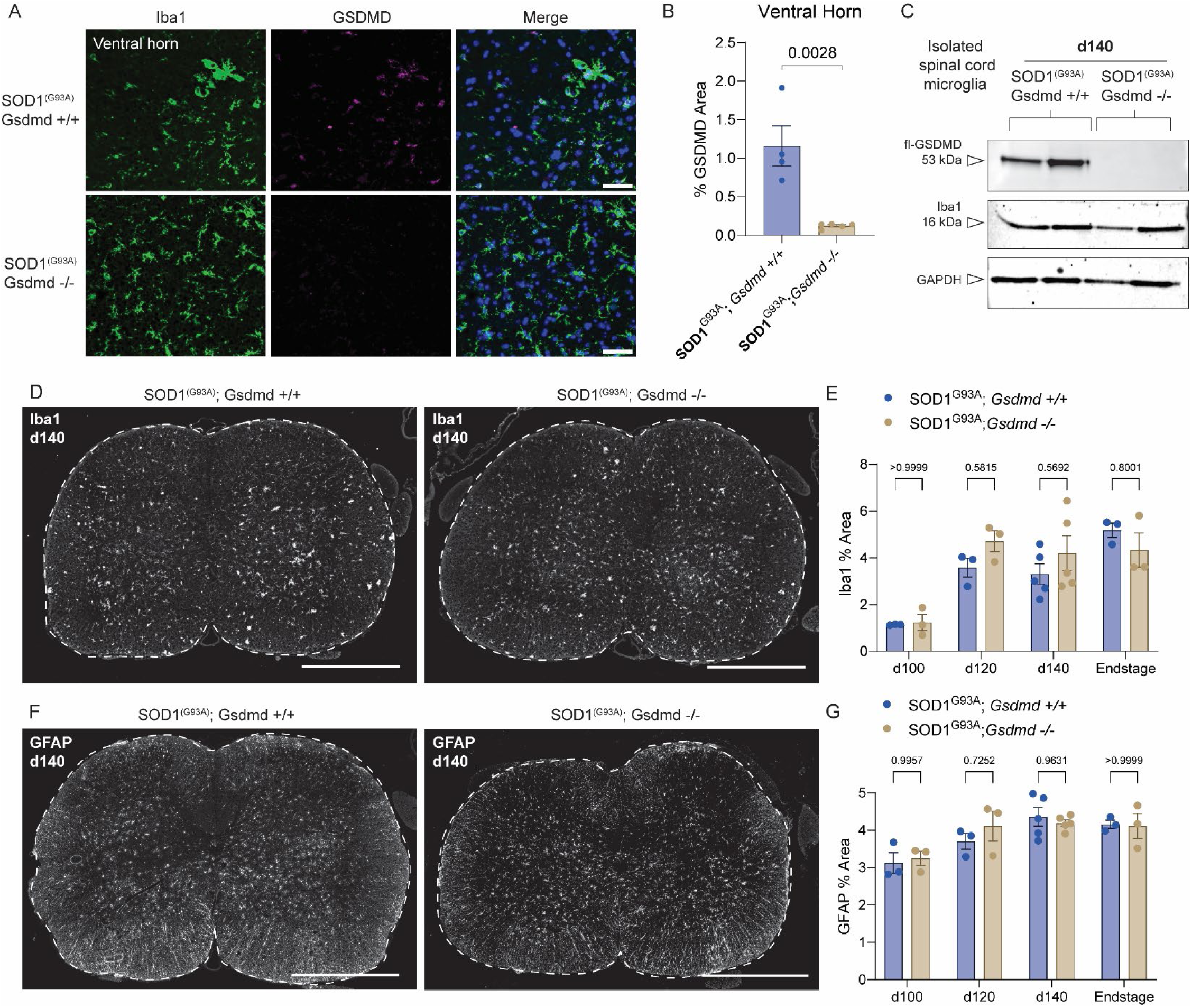
SOD1^G93A^; *Gsdmd* −/− mice do not display significant differences in microgliosis or astrogliosis compared to SOD1^G93A^; *Gsdmd* +/+ animals. (A) Representative images of the ventral horn of the lumbar spinal cord in B6 SOD1^G93A^; *Gsdmd*+/+ (upper panels) and SOD1^G93A^; *Gsdmd* −/− animals. Spinal cord sections were stained with microglia marker Iba1 (green) and GSDMD (magenta) with nuclei labeled with DAPI (cyan). Scale bar = 50um. (B) Quantification of percent GSDMD area normalized to total ventral horn area. Unpaired two-tailed Student’s t-test. N = 4 ventral horn ROI averaged from 4 SOD1^G93A^; *Gsdmd*+/+ animals, n = 4 ventral horn ROI averaged from 5 SOD1^G93A^; *Gsdmd*−/− animals. (C) Western blot of CD11b+ isolated cells (majority microglia) from the spinal cords of SOD1^G93A^; *Gsdmd*+/+ and SOD1^G93A^; *Gsdmd* −/− animals at day 140. (D) Representative images from SOD1^G93A^; *Gsdmd*+/+ (left) and SOD1^G93A^; *Gsdmd* −/− (right) lumbar spinal cords at 140 days stained with microglia marker Iba1. Scale bar = 500um. (E) Quantification of total Iba1+ area normalized to total spinal cord area at each timepoint (d100, d120, d140, and end-stage). (F) Representative images from SOD1^G93A^; *Gsdmd*+/+ and SOD1^G93A^; *Gsdmd* −/− lumbar spinal cords stained with astrocyte marker GFAP. (G) Quantification of total GFAP+ area normalized to total spinal cord area at each timepoint. Scale bar = 500um. (E,G) N = 3 spinal cord sections averaged for N = 3-5 animals per time point per genotype. Statistics in (E,G) performed using a Two-Way ANOVA with Šídák’s multiple comparisons test.

### 3.4 SOD1^G93A^; *Gsdmd−/−* mice display no change in disease onset or loss of grip strength

To establish whether loss of *Gsdmd* expression alters the rate or severity of disease progression in the SOD1-G93A model, we phenotyped male and female SOD1^G93A^; *Gsdmd+/+* and SOD1^G93A^; *Gsdmd−/−* mice starting from 60 days old until the full onset of hindlimb paralysis (∼d160-d180) (Figure 4 B-I). For our study, any animal that displayed transgene expression that was more or less than 30% of the overall mean transgene expression signal (AU) was excluded from the study. We tracked changes in individual animal body weight of SOD1^G93A^; *Gsdmd+/+* and SOD1^G93A^; *Gsdmd−/−* mice starting from 60 days old until mice succumbed to complete hindlimb paralysis and required euthanasia (Figure 4B-C). We found both male and female SOD1^G93A^; *Gsdmd+/+* and SOD1^G93A^; *Gsdmd−/−* mice had similar gains in overall body weight before the onset of disease resulted in decreasing body weight until survival endpoint (males, age F_(3.360, 84.24)_ = 42.67, p<0.0001; females, age F_(4.869, 134.2)_ = 42.25, p<0.0001) (Figure 4 B-C). However, we saw no effect of genotype on the percent change in body weight over time for either sex of SOD1^G93A^ transgenic animals (males, genotype F_(1, 26)_ = 2.490, p = 0.1266; females, genotype F_(1, 28)_ = 0.4930, p = 0.4884).

We first determined the age of disease onset, which is defined as the age at which animals reach peak body weight before the loss of motor neurons causes muscle to atrophy^34^. We found that female SOD1^G93A^; *Gsdmd+/+* mice reach peak body weight at 117.5 days of age, while female SOD1^G93A^; *Gsdmd−/−* mice reach peak body weight at 116.2 days (t=0.2961, df=28, p = 0.7693) (Figure 4 D). Male SOD1^G93A^; *Gsdmd+/+* mice reach peak weight at 112.3 days, while SOD1^G93A^; *Gsdmd−/−* males reach peak weight at 117.7 days (t=0.8917, df=24, p = 0.3814) (Figure 4 E). There were no significant sex differences in disease onset across the two genotypes. When sexes were combined, there was again no significant difference between SOD1^G93A^; *Gsdmd+/+* and SOD1^G93A^; *Gsdmd−/−* mice (t=0.5448, df=54, p = 0.5881) (Figure 4 F).

We next measured mouse grip strength across all four limbs every week from 80 days old to 160 days old (Figure 4 G-I). We found that both female and male SOD1^G93A^; *Gsdmd+/+* and SOD1^G93A^; *Gsdmd−/−* mice similarly lost grip strength with aging (females: age, F (8, 237) = 85.26, p<0.0001; males: age, F (8, 218) = 64.01, p<0.0001). However, there was no difference between genotypes for both females (genotype, F_(1, 237)_ = 0.0003160, p = 0.9858) and males (genotype, F_(1, 218)_ = 3.446, p = 0.0647) (Figure 4 G-H). By normalizing the grams of grip strength to the animal body weight at each age, we also combined the data across males and females and again while both sexes significantly lose grip strength, there is no effect of genotype (age, F_(8, 417)_ = 89.13, p<0.0001; genotype, F_(1, 417)_ = 0.0009878, p = 0.9749; interaction, F_(8, 417)_ = 0.4822, p=0.8689) (Figure 4 I).

We also phenotyped nontransgenic *Gsdmd*+/+ and *Gsdmd−/−* animals to determine any baseline changes in Iba1 reactivity, body weight, and grip strength (Supplemental Figure 4). Iba1+ area in the spinal cord for *Gsdmd* +/+ and *Gsdmd −/−* animals was not significantly different (t=0.3821, df=10, p= 0.7104) (Supplemental Figure 4 A,B). Male and female *Gsdmd* +/+ and *Gsdmd* −/− animals had slight differences in body weight as animals aged, with female *Gsdmd* −/− mice gaining slightly more wight over time than their *Gsdmd* +/+ counterparts (genotype, F_(1, 9)_ = 16.36, p = 0.0029) (Supplemental Figure 4 D). In contrast, *Gsdmd+/+* nTg males continued to gain weight over time, whereas male *Gsdmd−/−* animals plateaued at ∼100 days (genotype, F_(1, 18)_ = 6.165, p = 0.0231) (Supplemental Figure 4 G). Both male and female nontransgenic animals had steady grip strength over aging, with neither male nor female *Gsdmd*+/+ or *Gsdmd−/−* animals spontaneously losing grip strength over time with age (males, age F_(7, 141)_ = 0.5936, p =0.7603; females, age F_(7, 56)_ = 1.856, p = 0.0944) (Supplemental Figure 4 E, H). However, both male and female nontransgenic *Gsdmd−/−* mice had overall increased grip strength compared to *Gsdmd+/+* animals (males, genotype F_(1, 141)_ = 12.73, p = 0.0005; females, genotype F_(1, 56)_ = 18.70, p<0.0001). These baseline differences in bodyweight and grip strength did not impact the onset of paralysis in the SOD1^G93A^ transgenic animals, as we saw no differences between the two genotypes (Figure 4 D-F).

### 3.5 Loss of *Gsdmd* expression in SOD1^G93A^ mice does not alter microglia or astrocyte reactivity

To determine if loss of *Gsdmd* expression in our SOD1^G93A^; *Gsdmd−/−* transgenic mice would change the degree of microgliosis, we performed immunohistochemistry along a timecourse of disease (Figure 5 D). We analyzed spinal cords from SOD1^G93A^; *Gsdmd+/+* and SOD1^G93A^; *Gsdmd−/−* transgenic mice at non-symptomatic (d100), early symptomatic (d120), mid-stage disease (d140) and at end stage (full hindlimb paralysis). It has been well-established that the SOD1^G93A^ transgenic mice have significant increases in microglial numbers and reactivity as motor neurons degenerate in the spinal cord^35,36^. We analyzed the degree of microgliosis at each age across genotypes by quantifying the total Iba1 area over the entire spinal cord area (Figure 5 D,E). We found as SOD1^G93A^; *Gsdmd+/+* and SOD1^G93A^; *Gsdmd−/−* mice age, total Iba1+ area increases in the spinal cord (age, F_(3, 20)_ = 13.87, p<0.0001). However, there was no change in total Iba1+ area across genotypes (genotype, F_(1, 20)_ = 0.6295, p = 0.4369; interaction, F_(3, 20)_ = 1.198, p = 0.3359). We also analyzed the degree of astrogliosis by determining the extent of GFAP+ signal over the entire spinal cord area (Figure 5 F,G). As we found for the degree of microgliosis, while both genotypes displayed an increase in astrogliosis over time (age, F _(3, 20)_ = 7.919, p = 0.0011) there was no change in the amount of astrogliosis between SOD1^G93A^; *Gsdmd+/+* and SOD1^G93A^; *Gsdmd−/−* animals with aging (genotype, F_(1, 20)_ = 0.2105, p = 0.6514; interaction, F_(3, 20)_ = 0.5530, p = 0.6520).

### 3.6 GSDMD deficiency modestly accelerates mortality in SOD1^G93A^ mice

We next compared the survival rates over time between SOD1^G93A^; *Gsdmd−/−* and SOD1^G93A^; *Gsdmd+/+* mice. When segregated by sex, survival for female and male SOD1^G93A^; *Gsdmd−/−* was not significantly different than female or male SOD1^G93A^; *Gsdmd+/+* mice (females, Χ^2^ = 2.609, df= 1, p = 0.1062; males, Χ^2^ = 1.170, df= 1, p = 0.2793) (Figure 6 A-B). When data from both sexes of SOD1^G93A^; *Gsdmd−/−* mice were combined, survival was significantly worse in the SOD1^G93A^; *Gsdmd−/−* transgenic animals compared to SOD1^G93A^; *Gsdmd+/+* mice (Χ^2^ = 4.205, df= 1, p = 0.0403) (Figure 6 C). The median survival for combined sexes of SOD1^G93A^; *Gsdmd+/+* mice was 168 days while SOD1^G93A^; *Gsdmd−/−* animals have a median survival of 163 days.

**Figure 6.**
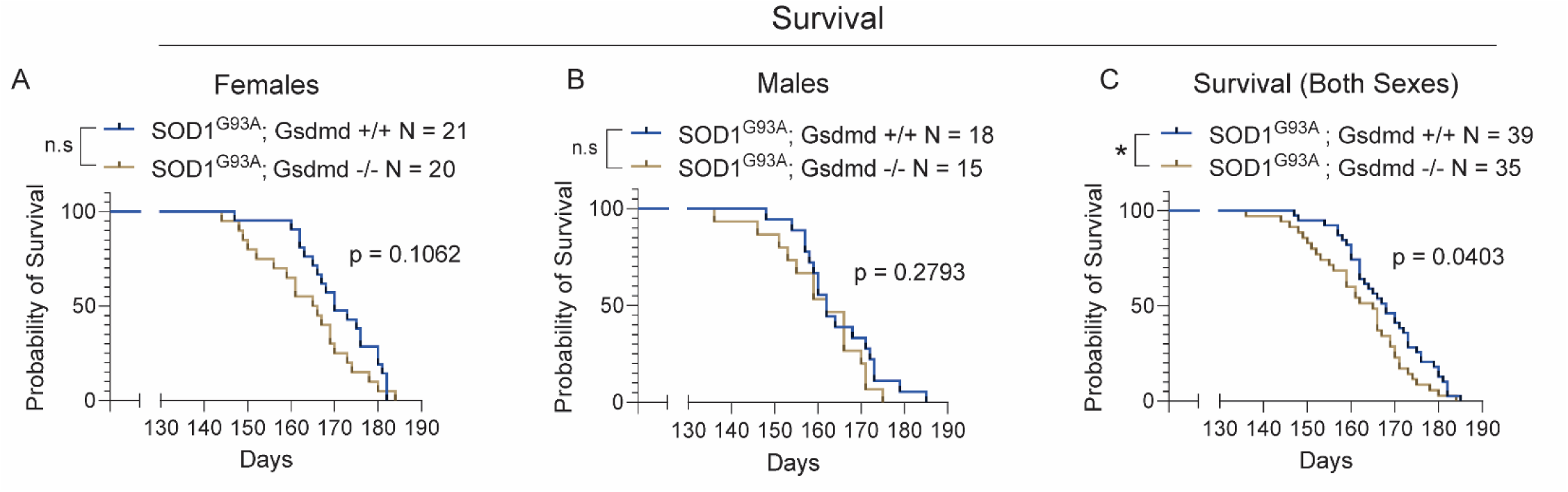
SOD1^G93A^; *Gsdmd* −/− mice show increased mortality compared to SOD1^G93A^; *Gsdmd* +/+ animals. Kaplan-Meier survival curves for female mice (A), male (B), and combined sexes (C) of SOD1^G93A^; *Gsdmd*+/+ and SOD1^G93A^; *Gsdmd* −/− animals. Numbers of mice: Females: SOD1^G93A^; *Gsdmd*+/+ mice, n=21; SOD1^G93A^; *Gsdmd*−/− mice, n=20, Males: SOD1^G93A^; *Gsdmd*+/+ mice, n=18; SOD1^G93A^; *Gsdmd*−/− mice, n=15; and combined sexes: SOD1^G93A^; *Gsdmd*+/+ mice, n=39; SOD1^G93A^*; Gsdmd*−/− mice, n=35. Statistical comparisons are performed using a Log-rank (Mantel-Cox) test.

We next determined any differences in the symptomatic phase of disease, which we defined as the number of days from peak body weight until death (Figure 7 A-C). There was no difference in the length of disease progression between male and female SOD1^G93A^; *Gsdmd+/+* and SOD1^G93A^; *Gsdmd−/−* mice (females, t=1.664, df=28, p = 0.1072; males, t=1.201, df=26, p = 0.2406). With sexes combined, the mean length of disease for SOD1^G93A^; *Gsdmd+/+* animals was 55.19 days, while the length for SOD1^G93A^; *Gsdmd−/−* mice was 47.04 days. This 8.15 day difference in disease length trended to where SOD1^G93A^; *Gsdmd−/−* have slightly shorter disease progression, indicating a slightly more severe disease course (t=1.990, df=56, p = 0.0514) (Figure 7 C).

**Figure 7.**
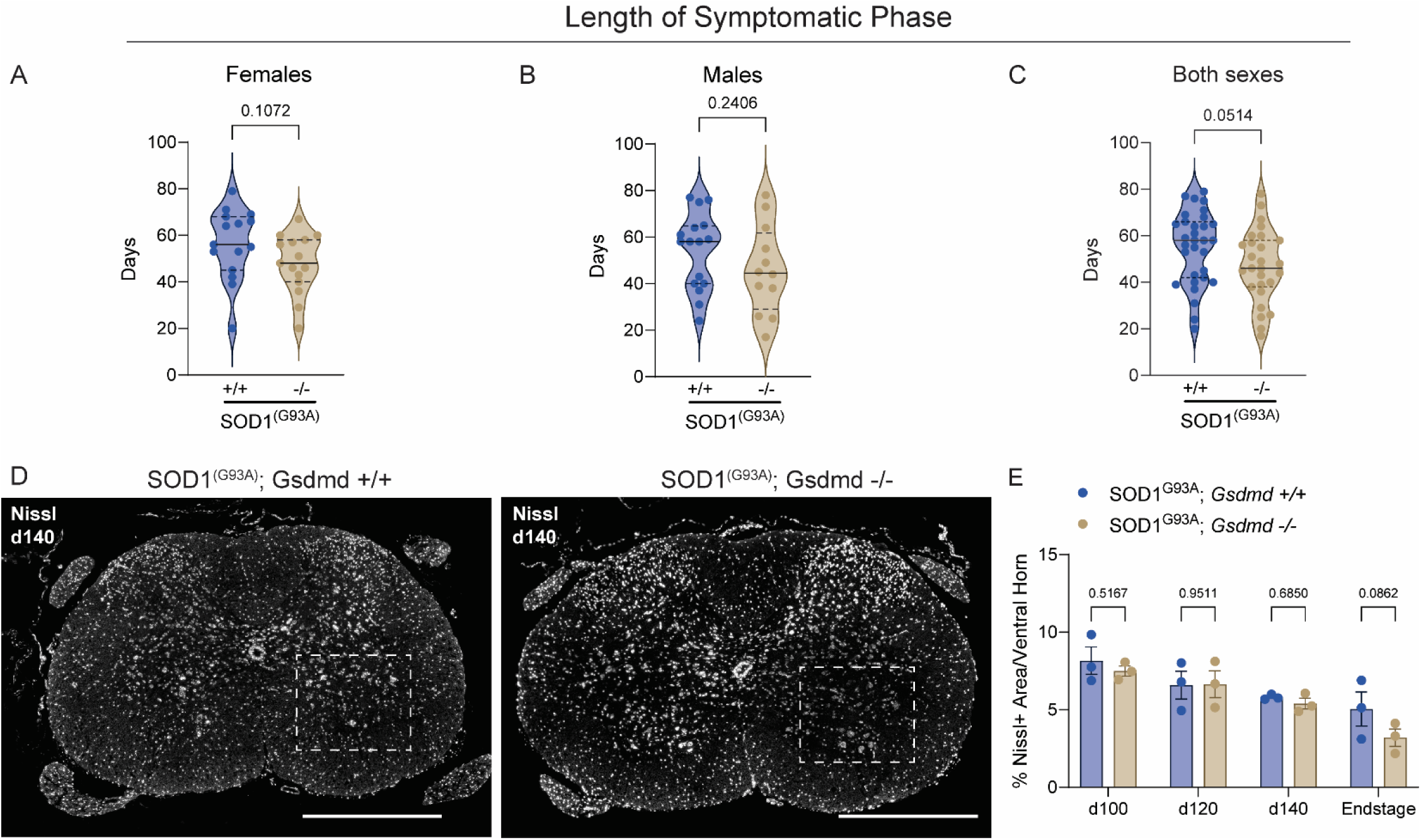
Gasdermin D deficiency causes minimal differences in symptomatic phase and motor neuron loss. (A-C) Quantification for the length of the symptomatic phase, defined as the number of days from peak body weight to euthanasia for females (A), males (B), and combined sexes (C). Numbers of mice: Females: SOD1^G93A^; *Gsdmd*+/+ mice, n=15; SOD1^G93A^; *Gsdmd*−/− mice, n=15, Males: SOD1^G93A^; *Gsdmd*+/+ mice, n=16; SOD1^G93A^; *Gsdmd*−/− mice, n=12; and combined sexes: SOD1^G93A^; *Gsdmd*+/+ mice, n=31; SOD1^G93A^; *Gsdmd*−/− mice, n=27. Statistics performed using an unpaired two-tailed Student’s t-test. (D) Representative images from SOD1^G93A^; *Gsdmd*+/+ and SOD1^G93A^; *Gsdmd* −/− lumbar spinal cords stained with Nissl. Scale bar = 500um. Dotted line represents ventral horn ROI used to quantify total Nissl+ neuron numbers at each timepoint. (E) Quantification of total Nissl+ neurons normalized to ventral horn area at each timepoint. N = 3 ventral horn ROI averaged per N = 3 animals per time point per genotype. Statistics performed using a Two-Way ANOVA with Šídák’s multiple comparisons test.

The hallmark of SOD1^G93A^-driven hindlimb paralysis is the loss of motor neurons in the spinal cord. We performed histology to determine if SOD1^G93A^; *Gsdmd−/−* animals have any change in the degree of neuron loss in the spinal cord with age. We performed Nissl staining to quantify the number of motor neurons in the ventral horn region of the lumbar spinal cord (Figure 7 D-E). We found that across both genotypes, there was a significant reduction in the total number of Nissl+ neurons with aging (age, F_(3, 16)_ = 9.698, p = 0.0007). However, this loss of neurons in the ventral horn was indistinguishable between SOD1^G93A^; *Gsdmd+/+* and SOD1^G93A^; *Gsdmd−/−* spinal cords (genotype, F_(1, 16)_ = 2.020, p = 0.1745; interaction, F_(3, 16)_ = 0.6459, p = 0.5968). At end-stage disease, there was a trend (p = 0.0662) of increased motor neuron loss in the SOD1^G93A^; *Gsdmd−/−* ventral horn compared to SOD1^G93A^; *Gsdmd+/+* animals (Figure 7 E), which correlates with the slight increase in the rate of mortality (Figure 6 C) and the trend towards more severe disease course (Figure 7 C) in SOD1^G93A^; *Gsdmd−/−* animals.

## 4. Discussion

In this study we sought interrogate the role of GSDMD in the disease pathology in the SOD1^G93A^ mouse model of ALS. We found that GSDMD is upregulated and activated (cleaved) in SOD1^G93A^ transgenic mice as they age, correlating with disease progression. In symptomatic SOD1^G93A^ mice, we also found concomitant increases in upstream activated caspase-1 and localization of GSDMD to Iba1+ microglial cells. Despite these dramatic changes in *Gsdmd* expression and activation in symptomatic mice, the loss of *Gsdmd* expression in a SOD1^G93A^ background did not ameliorate any facet of disease onset or progression *in vivo*. In fact, we found that the *Gsdmd* null SOD1^G93A^ mouse developed slightly more severe disease progression and worse overall survival. GSDMD has emerged as an attractive potential target for treatment in neurodegeneration: it is downstream of canonical inflammasome activation and found to be upregulated across several neurologic diseases^9,37^. However, the results from this study indicate that targeting GSDMD in the context of hereditary (mutant SOD1) ALS may not have a significant improvement on disease outcome. Our findings accord with earlier studies showing that chemical NLRP3 inhibition (MC905) did not alter disease progression in the SOD1 G93A mouse model^26^.

Increased reactivity and activation in resident microglia in the spinal cord is a hallmark of ALS^7^. Understanding how this microglial activation drives neurodegenerative disease severity remains an open area of investigation. Microglia reactivity can be a double-edged sword, where some inflammatory processes can confer neuroprotection while prolonged chronic inflammation may be harmful^38^. Understanding the contributions of microglial reactive state may depend on timing, for example in early versus late-stage disease, as well as spatial factors (i.e proximity to dying motor neurons). As we find null *Gsdmd* expression does not alter disease course in the SOD1 G93A model, the upregulation of *Gsdmd* in microglia during disease progression could suggest a potential protective or compensatory mechanism rather than a direct contribution to neurodegeneration. The trend towards increased mortality and motor neuron loss in SOD1^G93A^; *Gsdmd*−/− mice hints at the possibility that GSDMD could be involved in limiting microglia-driven neuroinflammation. For example, in the peripheral immune system GSDMD can negatively regulate neutrophil-mediated tissue damage by promoting neutrophil cell death^39^.

However as neurologic disease progresses, the chronic expression of *Gsdmd* in resident microglia may lead to chronic inflammatory cytokine release that may gradually exacerbate gliosis and neuron loss.

While we have focused on the role of GSDMD in resident spinal cord microglia in this model, we cannot rule out that other cell types express *Gsdmd* and may have important contributions to the severity of disease. For example, endothelial cells express *Gsdmd* in the spinal cord at baseline. A recent study elucidated a role for brain endothelial cells in driving blood brain barrier breakdown in the context of LPS-induced sepsis^40^. Loss of *Gsdmd* specifically in endothelial cells or microglia using conditional cell knockout lines may help further elucidate a potential harmful versus beneficial role of GSDMD activation throughout disease. Other cell types may also upregulate *Gsdmd* in a neurodegenerative context. For example, a study in the SOD1^G93A^ mouse model found that spinal cord astrocytes have increased NLRP3 expression^41^. Astrocytes could also upregulate *Gsdmd* expression and form GSDMD pores as disease worsens. Future studies are warranted to further dissect the role of GSDMD in different cellular contexts within the CNS and its potential dual roles in both neuroprotection and neurodegeneration. Understanding these dynamics could offer deeper insights into the complexities of ALS pathology and inform therapeutic strategies.

In summary, while we find GSDMD activation is associated with disease progression in the SOD1^G93A^ mouse model of ALS, global *Gsdmd* knockout does not significantly alter the disease course or neuroinflammatory response. These findings suggest targeting GSDMD in the context of mutant SOD1-driven ALS may be complex; the inflammasome-GSDMD axis may adopt opposing roles in early versus late-stage disease, as well as localization to different cell types. Future studies are warranted to understand cell-specific *Gsdmd* expression and activation throughout disease course in this model in order to develop significant therapeutic interventions targeting this molecular pathway.

## Authorship contribution statement

**Georgia Gunner**: Writing – original draft, Writing – review and editing, Conceptualization, Data curation, Formal analysis, Investigation, Methodology, Visualization. **Himanish Basu**: Conceptualization, Data curation, Methodology, Investigation, Writing – review and editing, Software, Validation. **Yueting Lu**: Data curation, Investigation, Formal analysis. **Matthew Bergstresser**: Data curation, Investigation, Formal analysis. **Dylan Neel**: Data curation, Conceptualization, Methodology, Writing – review and editing. **So Yoen Choi**: Data curation, Writing – review and editing. **Isaac Chiu:** Conceptualization, Supervision, Funding acquisition, Writing – original draft, Writing – review and editing, Resources, Project administration.

## Acknowledgments

We thank the Chiu lab members for helpful feedback. This work was supported by the Chan-Zuckerberg Initiative (CZI) Ben Barres Early Career Acceleration (ECA) award to I.M.C.; Target ALS grant to I.M.C.; NIH T32 AG000222 training grant to H.B. and S.Y.C., and NIH National Research Service Award (5F32AG084174-02) to G.G.

---

**Supplemental Figure 1:**
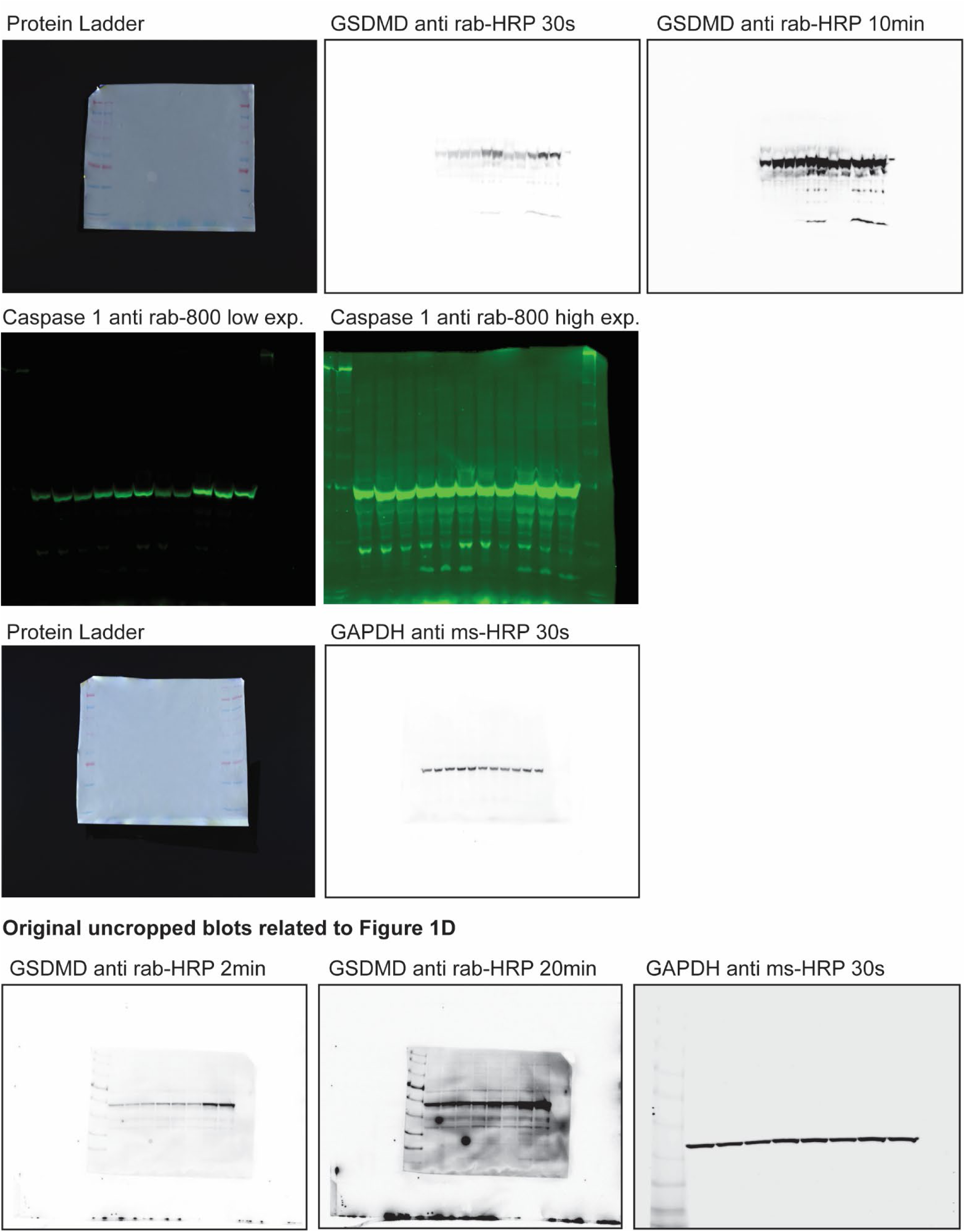
Original uncropped Western Blots related to Figure 1.

**Supplemental Figure 2.**
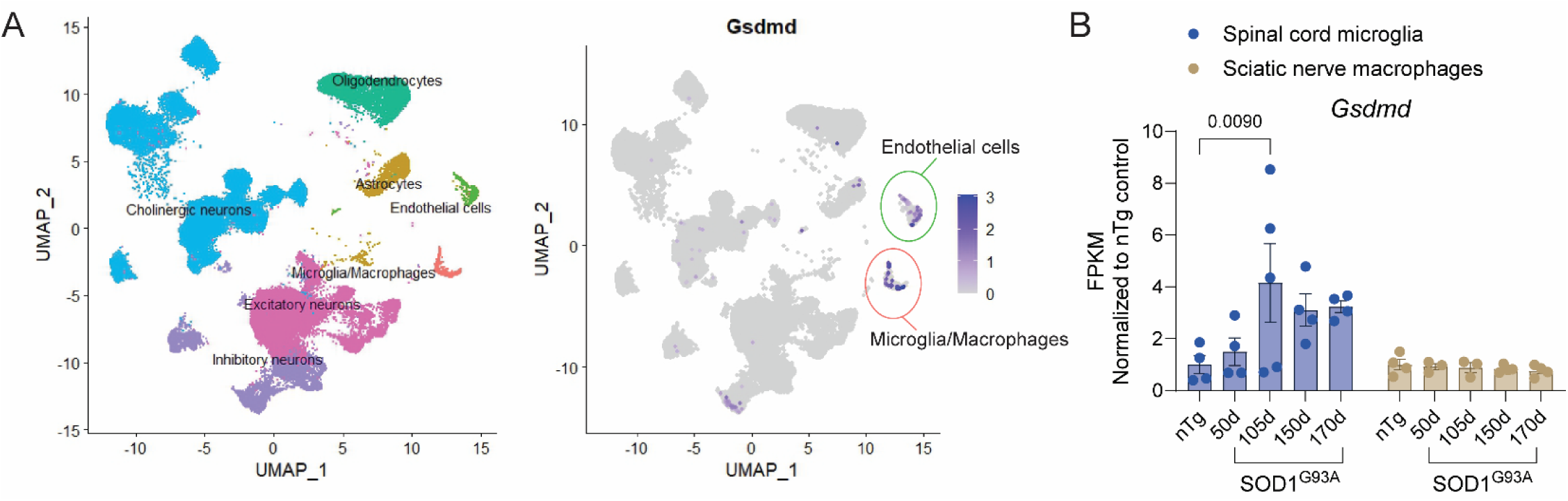
RNAseq data reveals expression of *Gsdmd* is specific to microglia and endothelial cells of the spinal cord and increases in SOD1^G93A^ microglia within the spinal cord as disease progresses. (A) Single nuceli RNA sequencing data from Blum et al 2021. UMAP of resident cell types in healthy, adult mouse spinal cord in left panel, expression of Gsdmd in right panel. (B) Bulk spinal cord microglia and sciatic nerve macrophage RNAseq data from a timecourse of SOD1^G93A^ transgenic mice from Choit et al 2020. Statistical analysis in (B) was performed using a Two-Way ANOVA with Šídák’s multiple comparisons test. N = 4-5 animals at each timepoint.

**Supplemental Figure 3:**
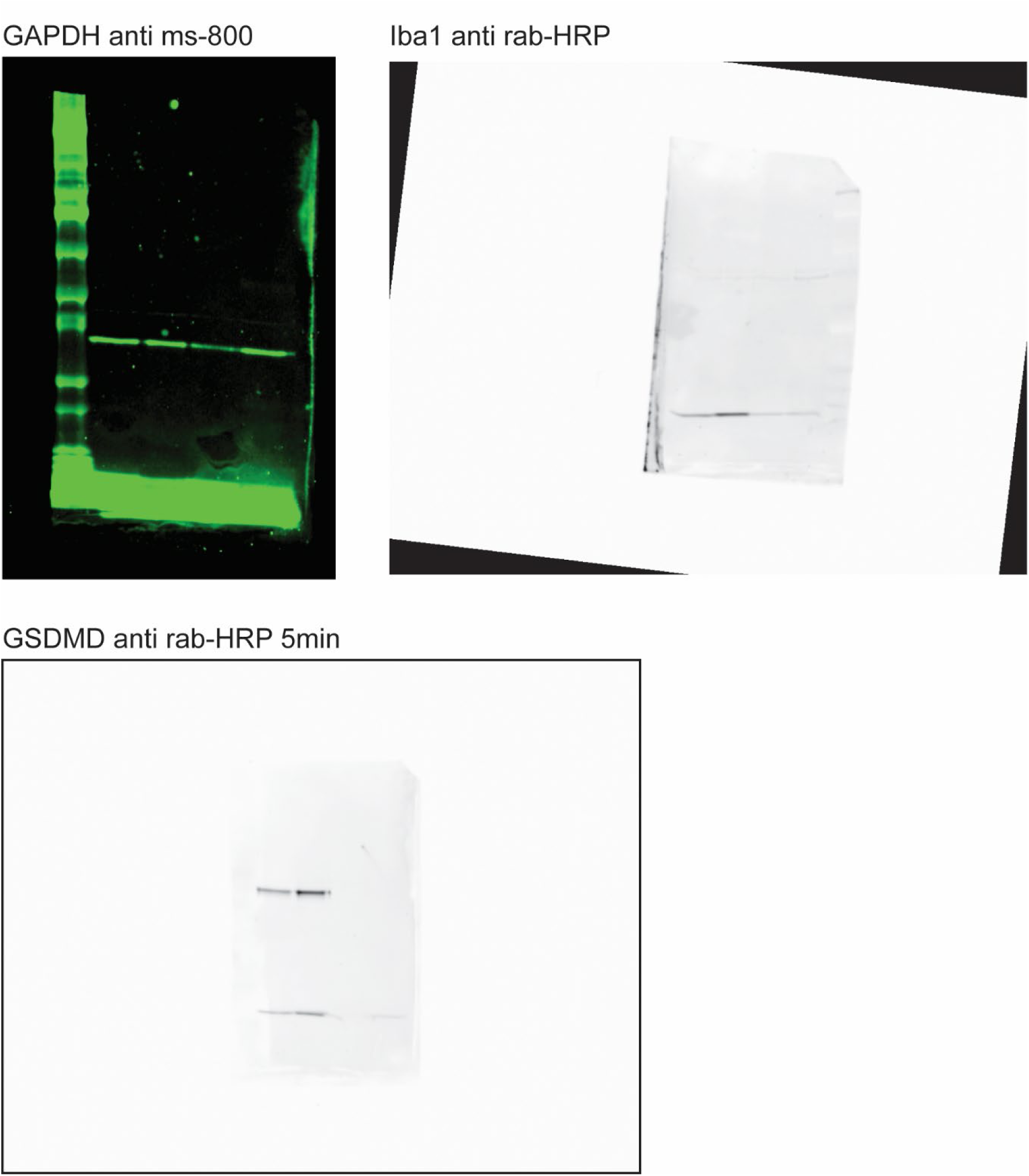
Original uncropped Western Blots related to Figure 3.

**Supplemental Figure 4:**
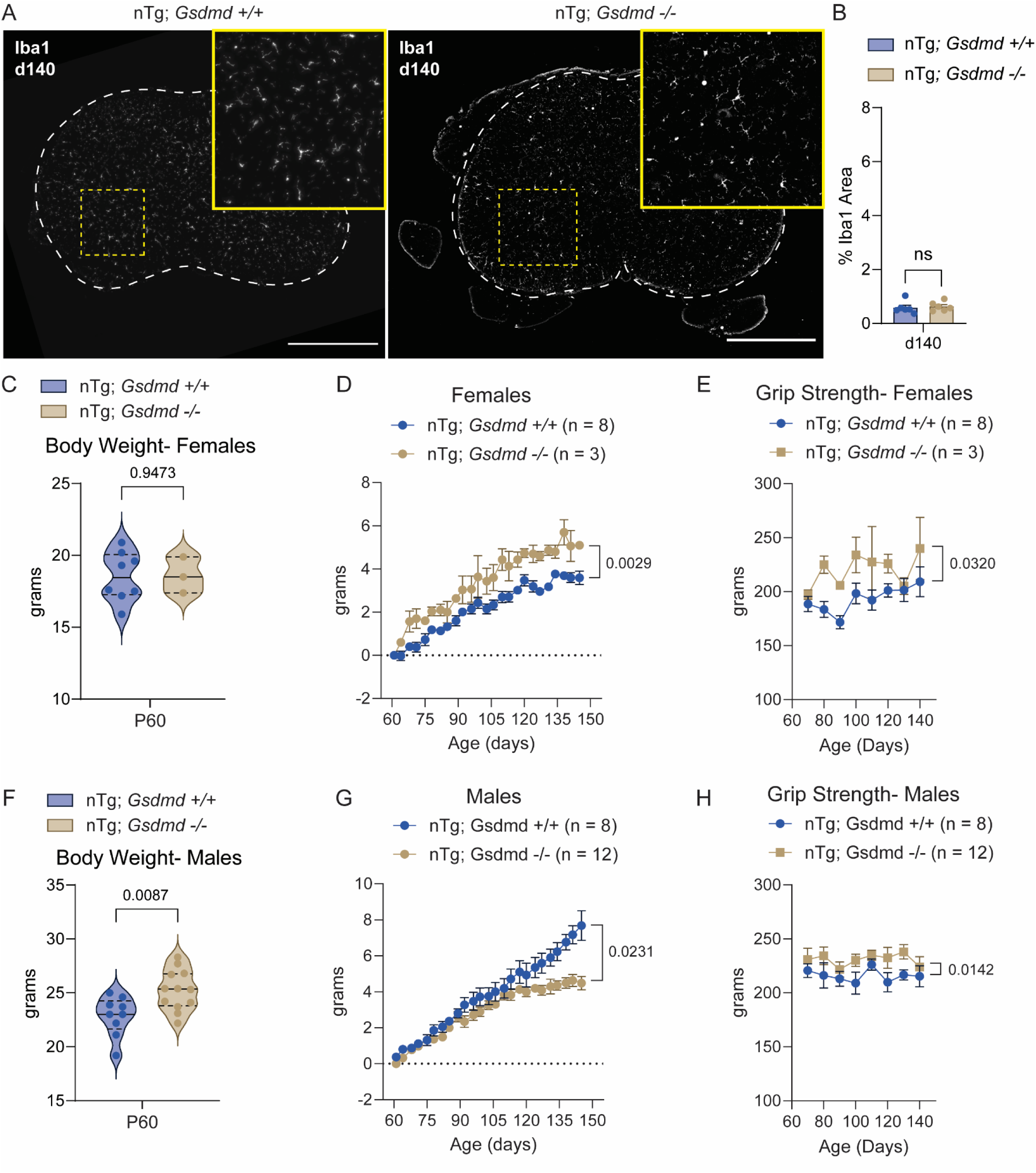
Nontransgenic *Gsdmd* −/− mice are grossly phenotypically similar to nontransgenic *Gsdmd* +/+ mice at baseline. (A) Representative images of lumbar spinal cord from d140 *Gsdmd*+/+ and *Gsdmd*−/− mice stained with microglia marker Iba1. Scale bar = 500um. (B) Quantification of percent total Iba1+ area normalized to total spinal cord section area. N = 3 spinal cord sections averaged per n = 6 *Gsdmd*+/+ and n= 6 *Gsdmd*−/− nontransgenic animals. Statistics performed using an unpaired two-tailed Student’s t-test. (C) Quantification for the average body weight of female nonstransgenic *Gsdmd*+/+ and *Gsdmd*−/− animals at 60 days of age. N = 8 *Gsdmd*+/+ and n = 3 *Gsdmd*−/− animals. Statistics performed using an unpaired two-tailed Student’s t-test. (D) Quantification for the change in body weight (grams) as female *Gsdmd*+/+ and *Gsdmd*−/− animals age. Statistics performed using a Two-Way ANOVA with Šídák’s multiple comparisons test. (E) Quantification for the change in grip strength over time for female nontransgenic animals. Statistics performed using a Two-Way ANOVA with Šídák’s multiple comparisons test. (F) Quantification for the average body weight of male nonstransgenic *Gsdmd*+/+ and *Gsdmd*−/− animals at 60 days of age. N = 9 *Gsdmd*+/+ and n = 12 *Gsdmd*−/− animals. Statistics performed using an unpaired two-tailed Student’s t-test. (G) Quantification for the change in body weight (grams) as male *Gsdmd*+/+ and *Gsdmd*−/− animals age. Statistics performed using a Two-Way ANOVA with Šídák’s multiple comparisons test. (H) Quantification for the change in grip strength over time for male nontransgenic animals. Statistics performed using a Two-Way ANOVA with Šídák’s multiple comparisons test

